# Functional MRI in awake dogs predicts suitability for assistance work

**DOI:** 10.1101/080325

**Authors:** Gregory S. Berns, Andrew M. Brooks, Mark Spivak, Kerinne Levy

## Abstract

The overall goal of this work was to measure the efficacy of fMRI for predicting whether a dog would be a successful service dog. The training and imaging were performed in 50 dogs entering advanced training at 17-21 months of age. FMRI responses were measured while each dog observed hand signals indicating either reward or no reward and given by both a familiar handler and a stranger. 49 dogs successfully completed fMRI training and scanning. Of these, 33 eventually completed service training and were matched with a person, while 10 were released for behavioral reasons. Using anatomically defined regions-of-interest in the ventral caudate, amygdala, and visual cortex, we developed a classifier based on the dogs' outcomes. We found that responses in the stranger condition were sufficient to develop an accurate brain-based classifier. On all data, the classifier had a positive predictive value of 96% with 10% false positives. The area under the receiver operating characteristic curve was 0.90 (0.79 with 4-fold cross-validation, P=0.02), indicating a significant diagnostic capability. Within the stranger condition, the differential response to [reward – no reward] in ventral caudate was positively correlated with a successful outcome, while the differential response in the amygdala was negatively correlated to outcome. These results show that successful service dogs transfer knowledge to strangers as indexed by ventral caudate activity without excessive arousal as measured in the amygdala.

## INTRODUCTION

The advent of awake fMRI in dogs has opened up numerous possibilities for decoding how the dog’s brain is organized (Berns et al., 2012; Andics et al., 2014; Jia et al., 2014). But like many of the early human fMRI studies, these nascent efforts have been plagued by small sample sizes. Some results have been replicated, increasing confidence in the technique (Berns et al., 2013; Dilks et al., 2015; Cuaya et al., 2016). At the same time, there have also been hints of substantial individual differences between dogs' brain responses that, like humans, relate to important aspects of behavior, temperament, and personality (Cook et al., 2014; Cook et al., 2016).

We have previously observed a temperament-dependent increase in neural activity in the ventral caudate when dogs are presented with a hand-signal associated with incipient receipt of food reward (Cook et al., 2014). Dogs who came from a service-dog training program were more likely to show a ventral caudate response to a hand signal when interacting with their owner/handler, while other dogs (e.g. from shelters and pets with no service-training) were more likely to show a caudate response to the signal when interacting with an unfamiliar person.

Are these differences a result of service-training, or might there be a neurobiological phenotype that predisposes a dog to becoming a good service-dog? If the latter, then it should be possible to identify good service-dogs before they complete service-training. Most dogs are not destined to be service dogs. Even with well-managed breeding programs, the success rate in training is typically 30-40%. By many estimates, the cost of training a service dog is $20,000 to $50,000. If dogs that are destined to fail training could be identified earlier, the average cost would decline. Thus, there is both a need to increase the number of service dogs and decrease the average cost by early identification.

There have been several studies of juvenile dogs, using behavioral tests and questionnaire ratings, that have examined the possibility of identifying high-potential working dogs. The earliest studies of puppy temperament indicated some trait consistency, but not to the extent of a diagnostic test (Scott and Fuller, 1965). Later studies showed that working dogs, on average, had different behavioral traits but these were not consistent enough to predict individual performance as adults (Goddard and Beilharz, 1986; Wilsson and Sundgren, 1997; Riemer et al., 2014). As dogs get older, their behavior becomes more consistent, achieving some level of stability between 6 and 12 months of age to the point where temperament questions and behavioral tests become modestly predictive of suitability for service work (Duffy and Serpell, 2012; Harvey et al., 2016). However, the variability of inter-rater agreement and test-retest reliability has raised questions about the utility of these approaches (Jones and Gosling, 2005; Fratkin et al., 2013).

To determine if a pattern of brain responses can predict completion of training and placement in a service job, we performed a prospective fMRI study of 50 dogs at the beginning of their service-training. We focused on three brain as potential biomarkers of success or failure: 1) ventral striatum for reward sensitivity (Schultz et al., 1997; Berns et al., 2013); 2) amygdala for arousal (Phelps, 2006); and 3) a region of temporal cortex previously shown to be responsive to faces (Dilks et al., 2015).

## METHODS

### Participants

All dogs participating in the study came from Canine Companions for Independence (CCI, Santa Rosa, CA). CCI-dogs undergo a rigorously controlled socialization process. After they are weaned, puppies are raised by a volunteer puppy-raiser until 17-21 months of age. Then, the dogs are returned to one of CCI’s training facilities for advanced training, which can take 6-9 months. Dogs that complete the training “graduate” into one of four roles: 1) service dog; 2) skilled companion; 3) facility dog; 4) hearing dog. Those who are unable to complete training, for either medical or behavioral reasons, are “released” and adopted, often by the puppy-raiser. The study was approved by both the Emory IACUC and UC Berkeley ACUC.

Between 11/2014 and 11/2015, 54 dogs were selected for participation in the MRI training program. The selection occurred within two weeks of beginning advanced training. Dogs were first assessed for absence of noise reactivity and then 2-3 dogs were randomly picked from each trainer’s string for participation in additional MRI training. Cohorts of 6-12 dogs were selected every 3 months, until the target of 50 was reached (5 dogs did not complete the training). One dog completed training, but did not successfully complete the MRI session. Four females were selected for breeding, and thus, no training outcome was obtained. Two dogs were released for medical reasons. This left a total of 43 dogs for which both MRI data and outcomes were obtained (Table 1). The dogs were predominately crosses of Labrador retrievers and golden retrievers, with a few purebred labs and goldens (Table 1).

**Table 1:** Demographics of dogs with both MRI data and outcomes. GLD=golden retriever; LAB=Labrador retriever; LGX=lab/golden cross.

|  | Breed |  |  | Total |
| --- | --- | --- | --- | --- |
|  | GLD | LAB | LGX |  |
| Female | 0 | 6 | 13 | 19 |
| Male | 1 | 2 | 21 | 24 |
| Total | 1 | 8 | 34 | 43 |

### Training

MRI-training took place on the CCI campus. A mock-MRI was constructed on site from discarded parts of a Siemens Trio. The mock-MRI included a patient table, bore, replicas of head and neck coils, and loudspeakers to play the noises from the scan sequences (Fig. 1). The training took approximately 10-15 minutes per dog, three days a week. All dogs were judged MRI-ready after two months.

The training program teaches the dogs to cooperatively enter the MRI scanner. The program is based on acclimatization to the MRI scanner noise, tight scanner enclosure, scanner steps, and operating vibrations and the shaping and ultimate chaining of several requisite behaviors. We also constructed a customized chin rest that facilitated comfort and proper positioning for the dogs and that adapted the apparatus for the uniqueness of the canine anatomy. Once the animals became confident and competent regarding all the preparatory steps – proven by completing a simulated MRI in the replica apparatus – we then performed live scans in the actual MRI.

**Fig. 1.**
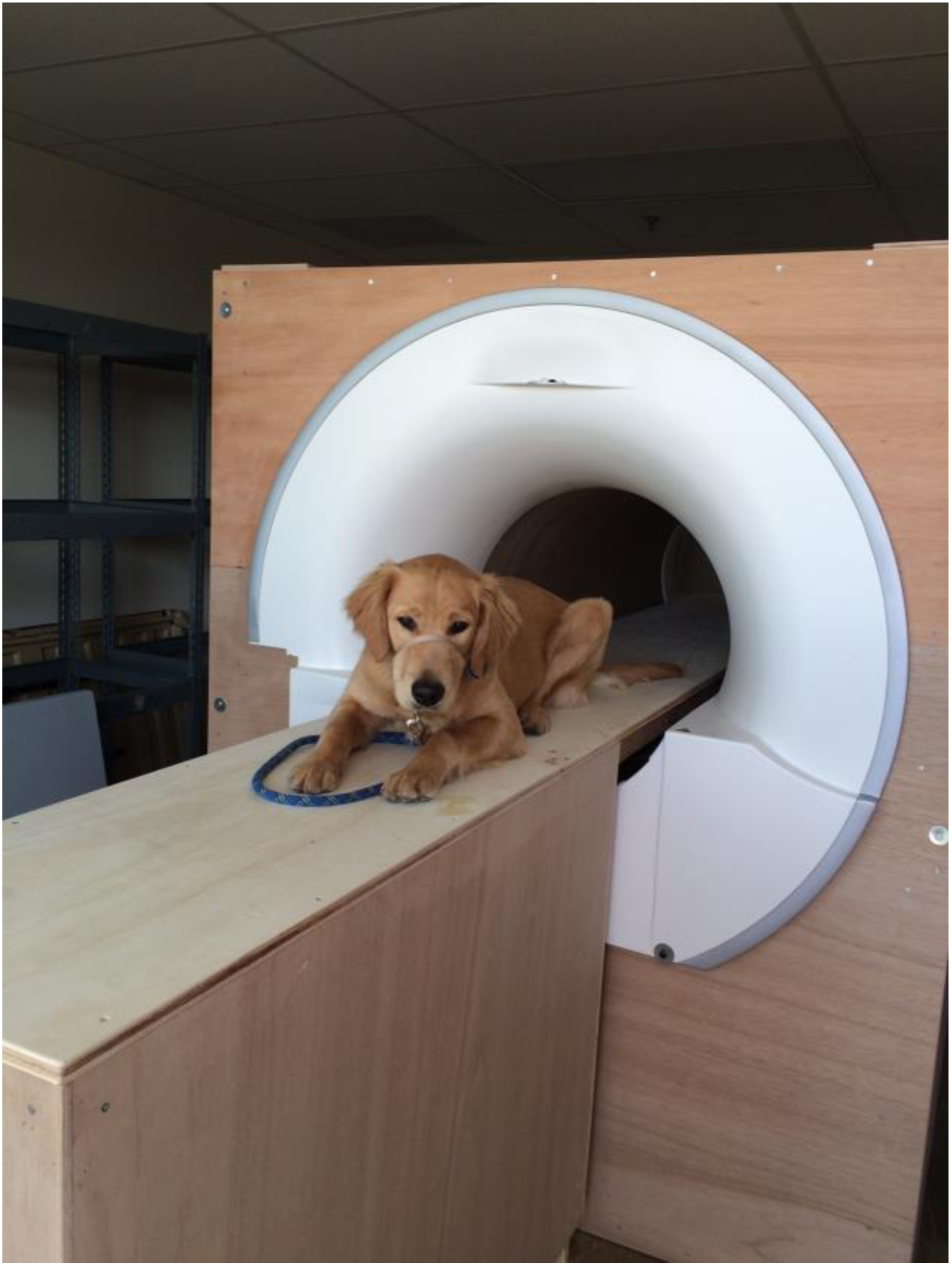
Mock scanner.

We compiled digitized audio recordings of the various scanner sequences. To aid in the necessary desensitization and acclimation to the scanner noises, when training, we played the recordings through a portable speaker placed inside the simulator.

Only positive reinforcement, in combination with behavioral shaping, conditioning and chaining, were used in the training process. First, dogs were trained to place their head and paws in the head coil. Next, they were trained to place their chin on a chin rest placed horizontally across the head coil and hold this position until a release signal. The length of the hold was gradually increased up to 30 s. When the dogs were able to do this consistently with no discernible head motion, they were next trained to do this wearing ear plugs and vet wrap, which were initially introduced to the animals apart from the coil simulator. Concurrent with the initial sequences of the training, recordings of the scanner noise were introduced at low volume. Once the animal became conditioned at a low volume, the volume was gradually increased. Recordings of the scanner noise were introduced at low volume while the dog remained stationary in the coil. Once the dog demonstrated relaxed behavior, the volume was gradually increased. When the dogs were comfortable wearing the ear plugs in the head coil with the scanner noise of approximately 90 dB, they were then trained to go into the MRI tube. Subsequently, the simulated head coil was placed inside the tube. After the dog was consistently holding her head still in this configuration, the entire apparatus was raised on a table to the height of the actual scanner patient table. The dogs were trained to walk up steps into the mock-MRI.

### Imaging

The MRI protocol is similar to that previously described (Berns et al., 2012; Berns et al., 2013). We have found that the neck coil that is standard on a Siemens Trio is ideally suited to scanning dogs' brains while in a crouch position. The chin rest is constructed from firm foam, and semicircles are cutout to match the shape of the dog’s muzzle from just the nose to the ramus of the mandible. We insert the dog’s custom chin rest within the inner diameter of the coil.

When performing an actual scan, immediately prior to the scan, we play audio recordings of the pertinent scan sequence through the scanner room speaker. As the dog settles in the scanner, we increase the recorded volume to match the decibel level of the actual scanner noise. While playing a continuous loop of the recording, once the sound level match we then begin the actual scan. The recordings are effective at minimizing the startle response that would otherwise result from the sudden onset of the real scan. Once the scan actual begins, we turn off the scanner recording.

First, a single sagittal plane image is acquired as a localizer, which lasts 3 s (SPGR sequence, slice thickness=4 mm, TR=9.2 ms, TE=4.16 ms, flip angle=40°, 256x256 matrix, FOV=220 mm). The localizer sound tends to be the most startling and unpleasant for the dogs. This is minimized by acquiring a single plane, but repeating it 3 times in case the dog startles at the onset. Because the chin rest centers the dog in the left-right direction, a single sagittal image is all that is necessary for planning the field-of-view for the subsequent scans.

After the localizer, a T2-weighted structural image is acquired with a turbo spin-echo sequence (28 2 mm slices, TR=3500 ms, TE=11 ms, flip angle = 131°, turbo factor =15, 128×128 matrix, 1.5 × 1.5 mm in-plane resolution), which lasts ~30 s. We use low-SAR and whisper mode to minimize acoustic noise. This sequence was optimized to yield good contrast between gray and white matter in the fastest time possible.

For functional scans, we use single-shot echo-planar imaging (EPI) to acquire volumes of 21 sequential 2.5 mm slices with a 20% gap (TE=25 ms, TR=1200 ms, flip angle=70°, 70x70 matrix, FOV=208 mm, 3 × 3 mm in-plane resolution). Slices are oriented transversely to the dog’s brain (coronal to the magnet because the dog is positioned 90° from the usual human orientation) with the phase-encoding direction right-to-left. Sequential scans are preferred to minimize between-plane offsets when the dog moves. The 20% slice gap minimizes crosstalk for sequential acquisitions. The right-left phase encoding minimizes ghost images from the neck that would otherwise overlap into the dog’s brain. TE is decreased slightly to minimize distortion. TR is as short as possible to acquire enough slices to cover the entire brain of most dogs while not so short as to significantly decrease signal.

### Experimental Design

The imaging protocol is an adaption of a previous experiment (Cook et al., 2014). During training, dogs are taught two hand signals: one representing a food treat (reward) and one representing no treat (no reward) (Fig. 2). Each hand signal is shown for 10 s, allowing enough time for the HRF to reach a peak before the dog moves to eat the treat. The key manipulation is who gives the signal: handler with whom the dog has been training or a stranger. There are four functional runs (each with 15 reward trials and 15 no-reward trials): 1) handler; 2) stranger; 3) stranger; 4) handler. The order of runs is counterbalanced in time to avoid confounding nonspecific effects due to repetition. Depending on the speed of the human, each run lasts 5-7 minutes, yielding 250-350 volumes/run, for a total 1000-1500 volumes. With time for acclimation and breaks between scans, the entire session lasts about 1 hour for each dog.

**Fig. 2.**
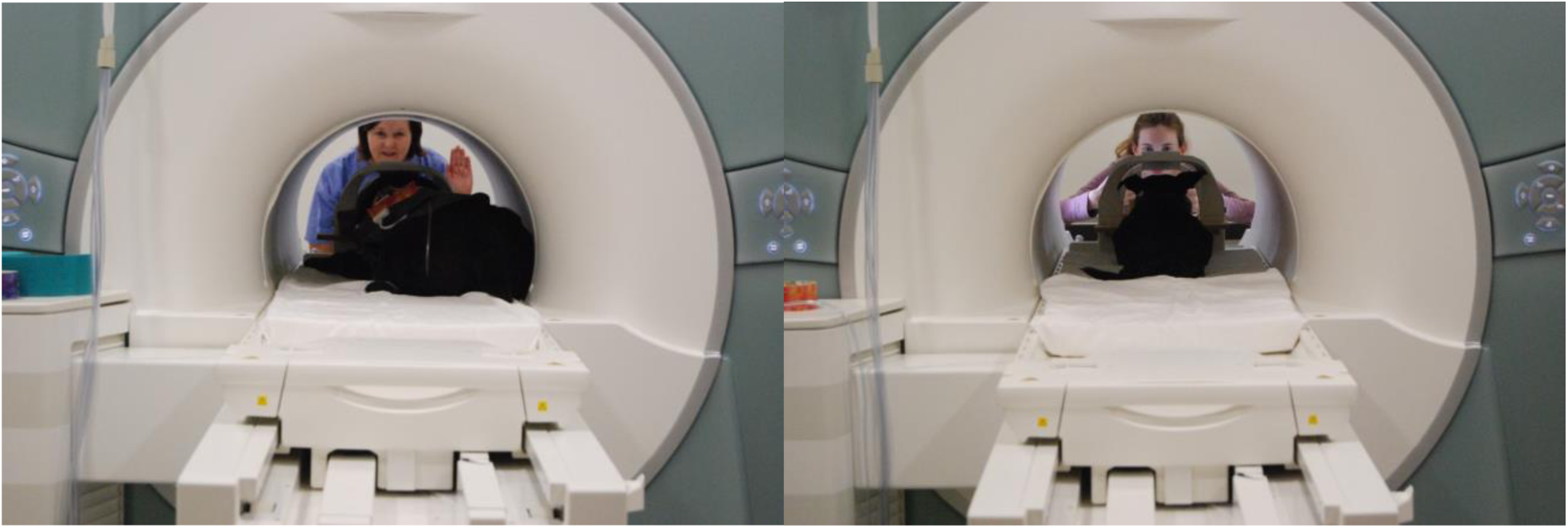
“Reward” hand signal (*left*) and “no reward” signal (*right*).

### Data Analysis

#### Preprocessing

Functional data was preprocessed using AFNI and its associated functions. DICOM files of the EPI runs were first converted to AFNI BRIK format using the to3d command. The EPI runs were then subjected to motion correction using 3dvolreg’s 6-parameter affine transformation, employing a two-pass method, where the first pass results in a crude alignment and the second pass a fine alignment. All volumes were aligned to a reference volume, which was manually chosen volume from the first run based on a visual inspection.

Because of the dog’s intermittent motion, censoring (removal) of volumes with artifacts is crucial. We used three separate methods to censor volumes with motion artifacts. First, 3dToutcount was used to output the fraction of outlier voxels for each volume. 3dToutcount defines outliers as those voxels whose signal intensity deviates from the median absolute deviation of the time series. Volumes with a fraction larger than 0.01 were censored from the statistical analysis. Second, 1d_tool.py was used to censor volumes based on the amount of estimated motion outputted from 3dvolreg. 1d_tool.py computes the Euclidean norm of the derivative of the rotation and translation parameters outputted from 3dvolreg. We then used a Euclidean norm cut-off of 1 to generate the censor file. Finally, we visually inspected the resulting time series with the censored volumes from 3dToutcount and 1d_tool.py, and censored any volumes that still showed obvious artifact. The majority of the censored volumes followed the consumption of the food reward.

The EPI images were then smoothed and normalized to %-signal change. Smoothing was applied using 3dmerge, with a 6 mm kernel at Full-Width Half-Maximum (FWHM). To convert signal intensity values to %-signal change, 3dcalc was used to subtract and then divide by the mean EPI image. These values were then converted to percentages by multiplying by 100.

#### General Linear Model (GLM)

For each subject, a GLM was estimated for each voxel using 3dDeconvolve. The task-related regressors included: 1) handler reward hand signal; 2) handler no-reward hand signal; 3) stranger reward hand signal; and 4) stranger no-reward hand signal. All events were specified as variable duration events using the dmUBLOCK function. To control for subject movement, the 6 motion regressors output from 3dvolreg were also included in the model. To account for differences between runs, a constant and linear drift term were included for each run. We generated 2 contrasts that depended on the source of the hand signals: 1) Handler: [reward – no reward]; and 2) Stranger: [reward – no reward].

#### Spatial Normalization

For each dog, three spatial transformations were computed: 1) rigid-body mean EPI to structural (6 dof); 2) affine structural to template (12 dof); and 3) diffeomorphic structural to template (Datta et al., 2012). These transformations were concatenated together and applied to individual contrasts obtained from the statistical model described above. The end result was a contrast image for each dog transformed into template space, allowing the computation of a group level statistic across all dogs. The transformations were computed using the software package, Advanced Normalization Tools (ANTs) (Avants et al., 2011).

#### Regions-of-interest (ROIs)

Based on previous experience, we defined 3 ROIs (Fig. 3): 1) bilateral ventral striatum (VS) at the location of maximal activation in previous studies with this paradigm in a different group of dogs (Berns et al., 2012; Berns et al., 2013; Cook et al., 2014); 2) bilateral amygdala (AMY); and 3) a region of visual cortex/temporal lobe previously identified to be face-selective in dogs, called dog face area (DFA) (Dilks et al., 2015). The amygdala ROI was based on the assumption that activity there would be related to arousal and could be relevant for predicting service dog success. The DFA was based on the possibility that differences in face-processing might also be relevant. The striatum and amygdala ROIs were spheres of 3 mm radius centered on these structures in template space. The DFA ROI was oblong and based on the location previously observed.

**Fig. 3.**
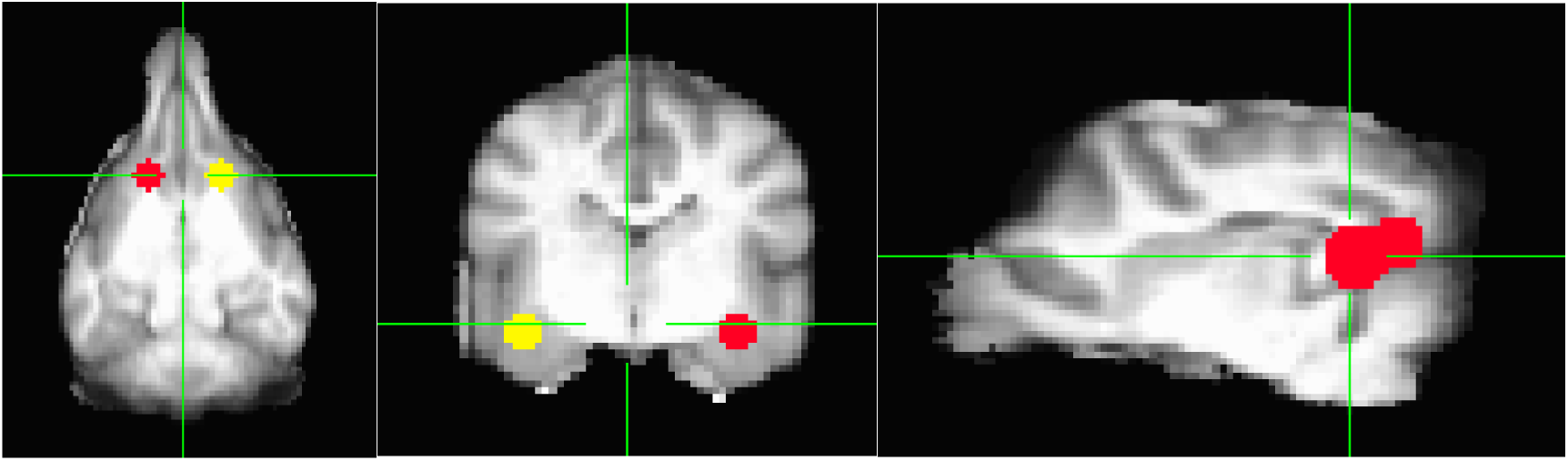
ROIs in: 1) ventral striatum; 2) amygdala; 3) dog face area of visual cortex.

#### Classifier

The primary goal of the project was to develop and test a brain-based classification algorithm that would predict successful graduation of service dogs. We used both the scikit-learn (http://scikit-learn.org) package in Python and glmfit function in Matlab to perform model-fitting, feature selection, and cross-validation. First, we performed feature selection by fitting both the ROI data and two demographic parameters: sex and responsiveness. The responsiveness score was assigned by the dogs' trainers mid-way through formal training based on the dog’s overall temperament, around the time of MRI-scanning. Values ranged from 1-6, with a higher score being more likely to succeed in the program and indicated a dog that is focused on and responsive to its handler. Responsiveness scores were assigned by handlers unaffiliated with the actual MRI training.

We tested several kernels including logistic, naïve Bayes, and linear support vector machine (SVM). We found that logistic regression consistently performed the best and so used that for feature selection. Because of the imbalance in dog outcomes, failures were weighted three times successes in the regression. First, we evaluated a behavioral-only model that consisted of sex and responsiveness score. This model is important because it provides a metric of performance that could be obtained without the need for brain-imaging. To be of practical use, a brain-based model should perform at least as well as behavior, and ideally, improve the performance of the classifier when combined with conventional behavioral measures like the responsiveness score. For feature selection, we tested the following models: 1) responsiveness; 2) sex + responsiveness; 3) 7 ROIs; 4) 4 ROIs; 5) sex + 4 ROIs; 6) responsiveness + 4 ROIs; 7) sex + responsiveness + 4 ROIs; 8) sex + responsiveness + 3 ROIs. The 7-ROI model consisted of the extracted contrast for [reward – no reward] from the VS, AMY, and DFA in both the handler and stranger runs. It additionally included the interaction between AMYxDFA for the stranger run on the possibility that there was an important interaction between face processing and amygdala activation. The 4-ROI models retained the terms for VS, AMY, DFA, and AMYxDFA from the stranger run only. And the 3-ROI model dropped the interaction term, retaining just VS, AMY, and DFA from the stranger run.

We calculated confusion matrices for each model with a threshold of 0.5, but for overall comparisons, we used the area-under-the curve (AUC) for the receiver operating characteristic (ROC), which plots true positive rate (TPR) vs. false positive rate (FPR) and thus does not depend on a particular threshold.

## RESULTS

The MRI-dogs had a high proportion of successes with 77% being placed in one of the assistance-dog categories (Table 2). Only 10 dogs were released for behavioral reasons. Although males slightly predominated in the failures, this was not statistically significant (Table 3).

**Table 2:** Outcomes of the dogs.

| Outcomes |  |
| --- | --- |
| Service Dog | 21 |
| Skilled Companion | 5 |
| Facility Dog | 4 |
| Hearing Dog | 1 |
| PTSD Dog | 2 |
| Behavioral Release | 10 |
| TOTAL | 43 |

**Table 3:**
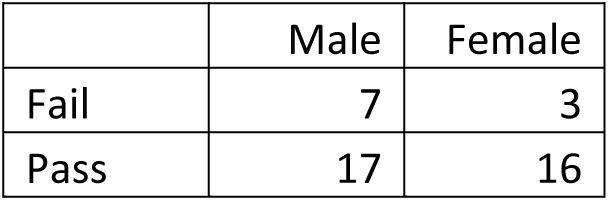
Distribution of successes and failures by sex. Sex was not a significant effect (Fisher’s exact test, P=0.47).

|  | Male | Female |
| --- | --- | --- |
| Fail | 7 | 3 |
| Pass | 17 | 16 |

As a check that the experiment was broadly consistent with previous imaging results, we first examined the whole-brain contrast for [reward – no reward] hand signals, averaged across all runs (both handler and stranger). This resulted in robust activation of the ventral striatum (Fig. 4), confirming the validity of the paradigm.

Both the behavior-only classifiers performed above chance. The sex+responsiveness model did very well, achieving an AUC=0.85 (Fig. 5, *blue*). The 7-ROI model achieved similar performance with an AUC=0.88. However, only the coefficients from the stranger run were significant, and so we dropped the features from the handler run, giving the sex+4-ROI model, which had an AUC=0.90 (Fig. 5, *green*). We then combined the sex+responsiveness features with the 4 ROIS (VS, AMY, DFA, AMYxDFA), which performed near perfectly with an AUC=0.96. Concerned that this model might overfit the data, the interaction term was dropped, giving the model of sex + responsiveness + 3 ROIs (from the stranger run) (Fig 5, *red*). This model had an AUC=0.95. The confusion matrices show the power of combining behavioral and brain measures (Table 4).

**Table 4:** Confusion matrices for different models. Threshold for predicted pass/fail=0.5.

|  | Behavior Only |  | Brain Only |  | Behavior + Brain |  |
| --- | --- | --- | --- | --- | --- | --- |
|  | Sex+Resp |  | Sex+VS+AMY+DFA+<br>AMYxDFA |  | Sex+Resp+VS+AMY+<br>DFA |  |
|  | Pred Fail | Pred Pass | Pred Fail | Pred Pass | Pred Fail | Pred Pass |
| True Fail | 8 | 2 | 9 | 1 | 9 | 1 |
| True Pass | 9 | 24 | 7 | 26 | 3 | 30 |
| NPV PPV | 47% | 92% | 56% | 96% | 75% | 97% |

**Fig. 4.**
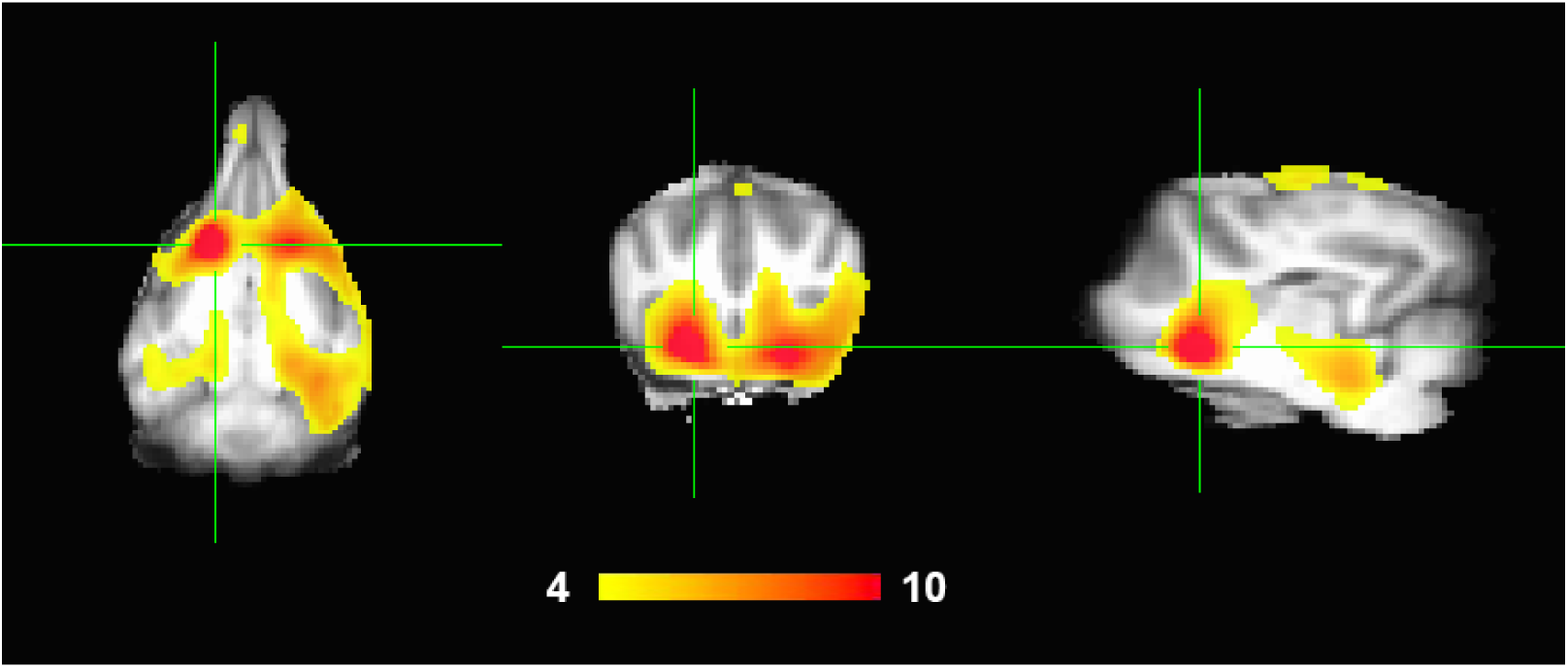
Average activation of [reward – no reward] hand signals, averaged across all conditions. Contrast is thresholded at p<10^−4^. Robust bilateral activation of ventral striatum is seen.

Importantly, even in the combined model, the coefficients for the VS and AMY were statistically significant (Table 5). <u>This indicates that the brain data added significant explanatory power even after accounting for sex and responsiveness</u>. Moreover, the signs of the coefficients are interpretable. Because female was coded as 1, and male as 0, the positive coefficient indicates that females had a higher chance of success, but only after accounting for the responsiveness score, which, as expected was also positively correlated with success. The differential response to [reward – no reward] was positively correlated in the ventral striatum and negatively correlated in the amygdala.

**Table 5:** Results of the full logistic model.

|  | coefficient | p |
| --- | --- | --- |
| intercept | -13.42 | 0.0025 |
| sex | 3.12 | 0.0285 |
| responsiveness | 2.81 | 0.0029 |
| VS str | 8.88 | 0.0034 |
| AMY str | -7.11 | 0.0032 |
| DFA str | -1.70 | 0.2222 |

**Fig. 5.**
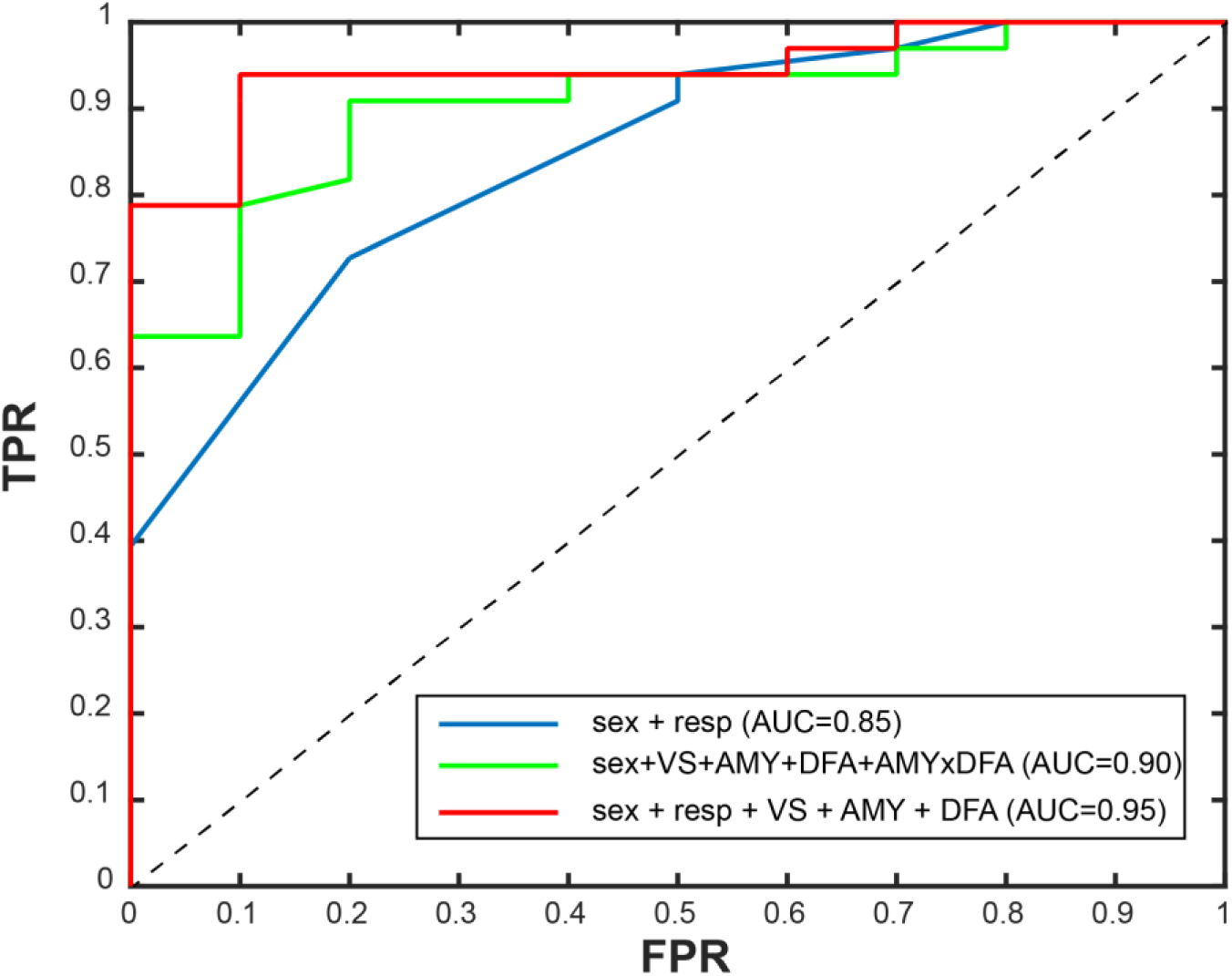
ROC plot of 3 classifier models. The combination of sex, behavior and 3 ROIs in the stranger-run performed the best.

Because these models were fit to all of the data, the performance represents a best-case scenario and would not be expected to do this well with new data (Fig. 6). To get a closer estimate of real-world performance, we performed the same classification of the final model using 10 iterations of stratified random shuffling, with a test size of 25%. Stratified random shuffling balances successes and failures for each split, which is important given our high class imbalance. We chose a 75/25 train/test split on each iteration due to the small sample size that would lead to high variance in accuracy estimates if smaller training sizes were used. As expected, the performance was not as good as the models fit to all of the data, but the AUC=0.79, which is still considered moderately good (Fig. 7).

**Fig. 6.**
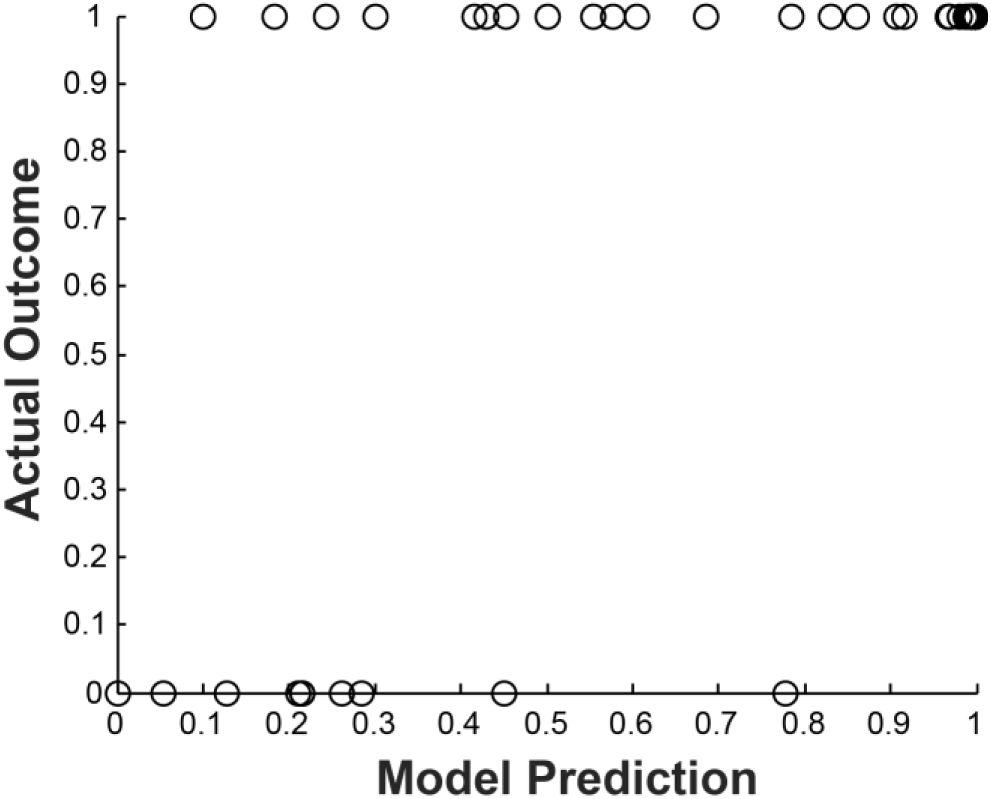
Fig. 6. Performance of the sex+brain model vs. actual outcome. 0=fail; 1=success.

## DISCUSSION

The primary goal of this study was to determine the efficacy of awake dog fMRI in predicting a dog’s suitability for assistance work. Using a paradigm in which the dog passively responded to hand signals indicating either incipient food reward or nothing, we found that the person giving the signals affected the dogs' brain responses. In particular, when a stranger gave the signals, instead of a familiar handler, we could use the differential activation in three regions-of-interest to predict the likelihood of a dog succeeding in assistance training.

**Fig. 7.**
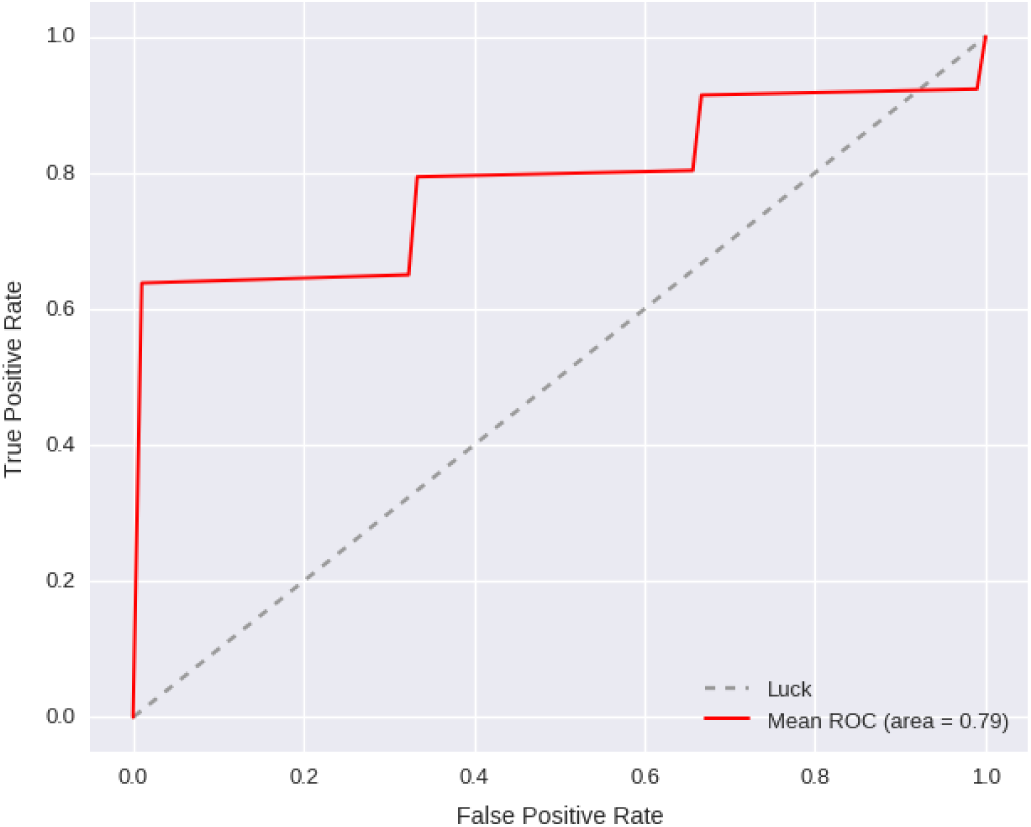
Estimate of real-world performance of the sex+ brain model using stratified random shuffling (P=0.02).

By itself, the brain-imaging model performed about the same as the sex+behavior model, with both achieving an AUC of 0.85-0.90. This alone may not warrant the use of brain imaging as a screening assessment for potential service dogs when common demographics and behavior tests do just as well. This level of predictive value is comparable to that reported in previous studies of behavioral tests of juvenile dogs (Harvey et al., 2016). However, combining brain activation with sex and behavior resulted in a significant improvement in predictive power, increasing the AUC to 0.95. Depending on the threshold chosen, this resulted in a PPV of up to 97% (Table 4). The brain data seems to be particularly valuable in decreasing the false negatives at any given threshold.

Although the results reported here represent the largest single fMRI study of dogs, the sample size is still considered small for predictive modeling. When the study was designed, we anticipated a 60% failure rate. Instead, we observed a 25% failure rate, which limits the generalizability of the model results. To mitigate the possibility of overfitting the model to the data, we performed cross-validation using stratified random shuffling. Stratification preserves the ratio of outcomes in each train/test iteration. With such a small sample size, we run the risk of noise dominating any individual iteration and so this type of validation is likely overly conservative, but this may counteract the optimistic performance obtained from the full dataset. The cross-validation suggested an AUC of 0.79 under real-world conditions, which is considered moderately good in terms of a diagnostic test.

Would such a procedure be worth the added cost of training and scanning dogs? The answer depends on the total cost to successfully place an assistance dog. Depending upon the vendor and the intended working role (e.g., psychiatric, mobility, hearing assistance), the cost to train a service dog may range from $20,000 to $50,000. The typical capital outlay for a nonprofit service dog organization that uses volunteer puppy raisers for the initial 12 to 18 months of the dog’s life is anywhere from $20,000 to $35,000. Thus, we can estimate the cost to raise and train a dog that does not get placed is still approximately $25,000. Conversely, there is also a cost for releasing a dog that might have become a service dog, but this may be relatively less if resources aren't expended to train her. Therefore, any cost savings would come from identifying dogs unlikely to succeed and releasing them from training as soon as possible.

Therefore, one should focus on the neuroimaging to identify dogs likely to fail. In a best case scenario, Table 4 suggests that 9 out of 10 dogs were correctly identified as failures (true negatives), and 7 were flagged incorrectly (false negatives). Clearly there is some cost to false negatives, but this is hard to estimate. Early identification of the 9 dogs could have saved approximately $225,000. If the cost of the MRI was $2000, then $86,000 would have been expended on all 43 dogs, for a net savings of $139,000. Would those dogs have been identified anyway? The sex+responsiveness model shows that 8 of them could have been, but at the cost of 2 more false negatives. However, the responsiveness score is assessed after the dogs are well into advanced training, after which they have already incurred costs. The true value of neuroimaging, then, can be realized primarily as an early screening tool, before dogs are selected for advanced training. For this to occur, dogs would need to begin MRI training at 12 months of age, and scanned at 15 months, when they would normally enter advanced training.

The higher than anticipated success rate of the dogs could be due to chance, although it is also possible that the MRI-training had a positive effect on the dogs. The training teaches self-control in the form of a highly disciplined ‘down-stay’ for minutes at a time in a noisy environment. It may be that this training generalizes to other skills necessary for an assistance dog.

Beyond the predictive value of the model, the relationship of brain activation in specific regions to outcome gives new insights into why some individuals are better assistance dogs. Even after controlling for sex and responsiveness, we found a significant relationship of outcome to activity in the ventral striatum and amygdala. We had hypothesized such relationships based on our previous findings that related striatal activity to cues signaling rewards like food and praise (Berns et al., 2012; Berns et al., 2013; Cook et al., 2014; Cook et al., 2016). Consequently, we designed the current experiment to measure the interaction between signals that designated reward and the source of the signal (familiar handler or a stranger). We found that the relative responses in striatum and amygdala in the handler conditions were not a significant predictor of outcome, but the responses in the stranger condition were. This was a surprise because we had hypothesized that the interaction would be the significant effect. Instead, the relevant responses could be isolated to the stranger condition.

Within the stranger condition, the differential response to [reward – no reward] in ventral striatum was positively correlated with a successful outcome, while the differential response in amygdala was negatively correlated to outcome. Considering first the striatal response, there is a vast literature linking ventral striatum, including nucleus accumbens, to positive expectation of reward (Schultz et al., 1997; Knutson et al., 2001; Montague and Berns, 2002; Berridge and Robinson, 2003; Ariely and Berns, 2010). It would seem that future assistance dogs can generalize the hand signals learned from their handler to a stranger. This ability to transfer knowledge learned from one person to another may be a key attribute of a good service dog.

The negative correlation with amygdala activation further clarifies the interpretation. Because amygdala activation can be seen in response to both positively and negatively valenced stimuli, its presence can be interpreted broadly as a measure of arousal. Here, arousal could occur due to either excitement or anxiety. But neither would be good for a service dog. Our results suggest that dogs who were more aroused by the stranger signals, as measured by the amygdala activation, were less likely to be placed as service dogs. Together, the results show that dogs successfully placed as service dogs transfer knowledge to strangers without excessive arousal.

## ACKNOWLEDGEMENTS

We are grateful to Erin Rich, Amy Kinsella, and Shelley Lampman for their skill in training the dogs. Ben Ingles and Rick Redfern of the Henry H. Wheeler, Jr. Brain Imaging Center were a tremendous help in scanning the dogs at UC Berkeley. Marian Scopa coordinated the logistics of the project.

This material is based upon work supported by the Army Contracting Command and DARPA under Contract No. W911NF-14-C-0094. Any opinions, findings and conclusions or recommendations expressed in this material are those of the authors and do not necessarily reflect the views of the Army Contracting Command and DARPA. G.B. and M.S. own equity in Dog Star Technologies and developed technology used in some of the research described in this paper. The terms of this arrangement have been reviewed and approved by Emory University in accordance with its conflict of interest policies.

### AUTHOR CONTRIBUTIONS

The experiment was designed by G.B., A.B., and M.S. Training was supervised by K.L. All authors participated in scanning, and K.L. collected additional behavioral data. Analysis was performed by G.B. and A.B. The manuscript was written by G.B and edited by all authors.

